# Influence of rotational nucleosome positioning on transcription start site selection in animals promoters

**DOI:** 10.1101/064113

**Authors:** René Dreos, Giovanna Ambrosini, Philipp Bucher

## Abstract

The recruitment of RNA-Pol-II to the transcription start site (TSS) is an important step in gene regulation in all organisms. Core promoter elements (CPE) are conserved sequence motifs that guide Pol-II to the TSS by interacting with specific transcription factors (TFs). However, only a minority of animal promoters contains CPEs. It is still unknown how Pol-II selects the TSS in their absence. Here we present a comparative analysis of promoters’ sequence composition and chromatin architecture in five eukaryotic model organisms, which shows the presence of common and unique DNA-encoded features used to organize chromatin. Analysis of Pol-II initiation patterns uncovers that, in the absence of certain CPEs, there is a strong correlation between the spread of initiation and the intensity of the 10 bp periodic signal in the nearest downstream nucleosome. Moreover, promoters’ primary and secondary initiation sites show a characteristic 10 bp periodicity in the absence of CPEs. We also show that DNA natural variants in the region immediately downstream the TSS are able to affect both the nucleosome-DNA affinity and Pol-II initiation pattern. These findings support the notion that, in addition to CPEs mediated selection, sequence-induced nucleosome positioning could be a common and conserved mechanism of TSS selection in animals.

**Author Summary:** Gene transcription is a complex process that starts with the recruitment and positioning of Pol-II enzyme at the transcription start site (TSS). Specific promoter sequences, known as core promoter elements (CPEs) facilitate this process. Surprisingly, only a fraction of promoters contain them. It is still unknown how Pol-II choses the start site in their absence. Arecently proposed alternative mechanism implicates positioned nucleosomes in the TSS selection. Here, we provide new evidence of the existence of such mechanism with a comparative analysis of promoter’s features across the animal kingdom. We analysed the promoter’s DNA sequence composition in 5 organisms and found conserved and unique consensus sequences used to organize chromatin in the region of the first nucleosome downstream the TSS (N+1). Moreover, we found that all organisms show a strong correlation between the spread of Pol-II initiation and the strength of the DNA-encoded signal in the N+1 region. A detailed analysis of Pol-II initiation sites reveals also the presence of a 10 bp periodicity that is correlated with the intensity of the DNA signal in the N+1 region. Importantly, we report that genetic variants that alter the DNA-nucleosome affinity in that region alter Pol-II initiation spread as well.

## Introduction

An essential step in gene regulation is the recruitment of RNA-Pol-II (Pol-II) to the transcription start sites (TSS) at gene promoters [1–3]. This is often facilitated by the presence of conserved sequence motifs known as core promoter elements (CPEs), which are found at a fixed or nearly fixed distance from the TSS [4, 5]. Among them, the TATA-box, located 25-30 base-pairs (bp) upstream of the TSS, and the Initiator (Inr), located at the TSS, are the best known and most widely conserved CPEs among species [6, 7]. The TATA-box is bound by general transcription factors (TFs) that guide and anchor Pol-II to the TSS [8]. As a consequence, promoters with a TATA-box are generally characterized by a focused, almost to the single base, start site [9, 10].

In spite of the CPE’s demonstrated capability to select a TSS with high precision, only a minority of promoters have a CPE (in human 10% a TATA-box, 30% an Inr motif) [11]. A central question in gene expression is how Pol-II selects the TSS in their absence [12, 13]. It has been shown that nucleosomes in promoter regions can regulate gene expression via TF binding site occlusion [14] but their role in TSS selection by Pol-II remains unclear. Promoters have a remarkably conserved chromatin architecture consisting of a nucleosome free region that spans 100-150 bp upstream the TSS followed by a well-positioned nucleosome (+1 nucleosome) [15, 16]. This general conformation can be altered by diverse factors. Contrary to intuition, so called broad promoters with dispersed initiation sites have the most focused and regular nucleosome architecture whereas narrow promoters (also referred as peak promoters) have less organized nucleosomes [17] and an atypical chromatin architecture [18]. In zebra fish, the chromatin architecture of the same promoter has been shown to change from one developmental stage to another [19] but there again, the conformation with the more structured nucleosome architecture shows a broader initiation site pattern. In mammals, promoters have traditionally been classified according to the presence or absence of CpG islands (CGI), 500-1000 bp long regions enriched in C+G [20–22]. CGI-promoters are often TATA-box depleted [23], have broad TSS [9], exhibit characteristic histone marks [24] and have a precisely positioned +1 nucleosome which is present even when the promoter is not transcribed [25]. In essence, CGI-promoters resemble the broad promoters described in other species and thus may not be considered a separate class.

An open question in gene regulation is whether the chromatin at promoters is organized by sequence-intrinsic features or indirectly by the transcription machinery occupying the nucleosome-free region and thereby forcing the nucleosome to bind to the nearest free space downstream the TSS. On a genome level, two types of sequence features have been reported to participate in nucleosome positioning: dinucleotide periodicity and base composition [26]. A theoretical model suggests that the same dinucleotide repeated at 10 bp intervals leads to intrinsic curvature that favors the wrapping of the DNA around the histone octamer [26, 27]. This model theorizes that the periodic dinucleotide always occurs with the same orientation relative to the histone-octamer surface, for instance having the major groove facing outwards, and implies a rotational positioning of the nucleosome. Some authors have identified WW (W for A or T) and SS (S for C or G) dinucleotides in counter-phase as major contributors of rotational positioning [28, 29], others emphasized the importance of RR (R for A or G) and YY (Y for C or T) motifs [27]. DNA base composition can also affect nucleosome positioning. Highly AT-rich sequences, in particular poly(dA:dT) tracts, strongly disfavor nucleosome formation [30, 31], whereas G+C rich sequence tend to have high nucleosome occupancy [32, 33]. Unlike dinucleotide periodicity, sequence composition can position nucleosomes in a narrow DNA region without specific preference for rotational setting, a condition termed translational positioning.

As said before, the role of sequence-intrinsic features in chromatin organization around promoters remains a matter of debate [34]. Zhang and colleagues concluded that its positioning is the result of Pol-II binding to the DNA [35]. Recent studies done in yeast have shown that chromatin remodelers play an important role in organizing chromatin both at a genome [36] and promoter level [37] and that they act synergistically with DNA sequences [38]. Others have reported the presence of nucleosome-favoring sequences in yeast promoters [27, 39–41], with a high correlation between *in-vitro* and *in-vivo* nucleosome organization in these regions [42, 43]. Recently, a 10 bp periodic signal has been observed in cumulative WW frequency plots of promoters sequences aligned with respect to the major TSS as defined by CAGE [44]. A similar WW periodicity can be seen in WW heat map plots published in [19]. The phasing of WW periodicity with the TSS is the first indication that the rotational setting of the DNA in the +1 nucleosome is guiding the TSS selection process.

In this paper, we investigate the molecular mechanisms of TSS selection by jointly analyzing experimentally determined chromatin architectures, DNA-encoded nucleosome signals, Pol-II initiation site patterns and natural genetic variation in promoters stratified by the presence or absence of specific CPEs and/or the breadth of the initiation patterns. The analysis on five model organisms(*Homo sapiens, Mus musculus, Danio rerio, Drosophila melanogaster* and *Caenorhabditis elegans*)confirms that different species have an overall similar chromatin organization with nevertheless some noteworthy species-specific differences. All five organisms have sequence-intrinsic nucleosome-positioning signals that are predictive of *in-vivo* nucleosome organization, but only in promoters that lack TATA-boxes. Additionally, we show that broad promoters associated with strong sequence-encoded nucleosome +1 have 10 bp periodic initiation patterns. By analyzing the effects of genetic variants on promoter initiation site patterns and dinucleotide periodicity, we provide genetic evidence that rotational nucleosome positioning is mechanistically involved in TSS selection.

## Results

### Promoters have DNA rotational properties that influence *in-vivo* nucleosome organisation and are affected by species-specific biases in DNA composition

To verify that DNA sequences around animal promoters had rotational nucleosome-positioning properties and that the 10-11 bp was the prevailing frequency, 1 kb regions on each side of *H. sapiens, M. musculus, D. rerio, D. melanogaster* and *C. elegans* TSSs were scanned for the presence of periodic signals of any length for each individual WW, SS, YY, or RR dinucleotide (S1 Fig). Confirming our expectations, all organisms showed a peak in signal intensity for periods of 10-11 bp (S2 Fig) that are typical of nucleosomal DNA with a minimum in correspondence of the nucleosome free region and a maximum in the N+1 region (S3 Fig). To further study the rotational properties of single promoters sequences and their effect on chromatin conformation, the strength of 10.3 bp periodic signals for each dinucleotide was evaluated in each promoter and compared to their *in-vivo* nucleosome maps. As expected, the WW dinucleotide (or SS for *D. melanogaster)* showed the highest correlation with *in-vivo* nucleosome signals (Figs 1A and S4). In *H. sapiens*, about one third of promoters (top promoters of Fig 1A) had low WW periodicity upstream the TSS and a peak in periodicity immediately downstream. This was reflected in the chromatin organisation with a clear nucleosome free region (NFR) and a focused N+1. As expected, this group of promoters was also depleted of TATA-box and enriched in CpG islands. Approximately 25% of promoters showed an opposite signal, with a peak upstream the TSS and a valley downstream (promoters at the bottom of Fig 1A). They were characterized with a less pronounced NFR, a broader N+1, an enrichment in TATA-box and depletion in CpG islands.

**Fig. 1.**
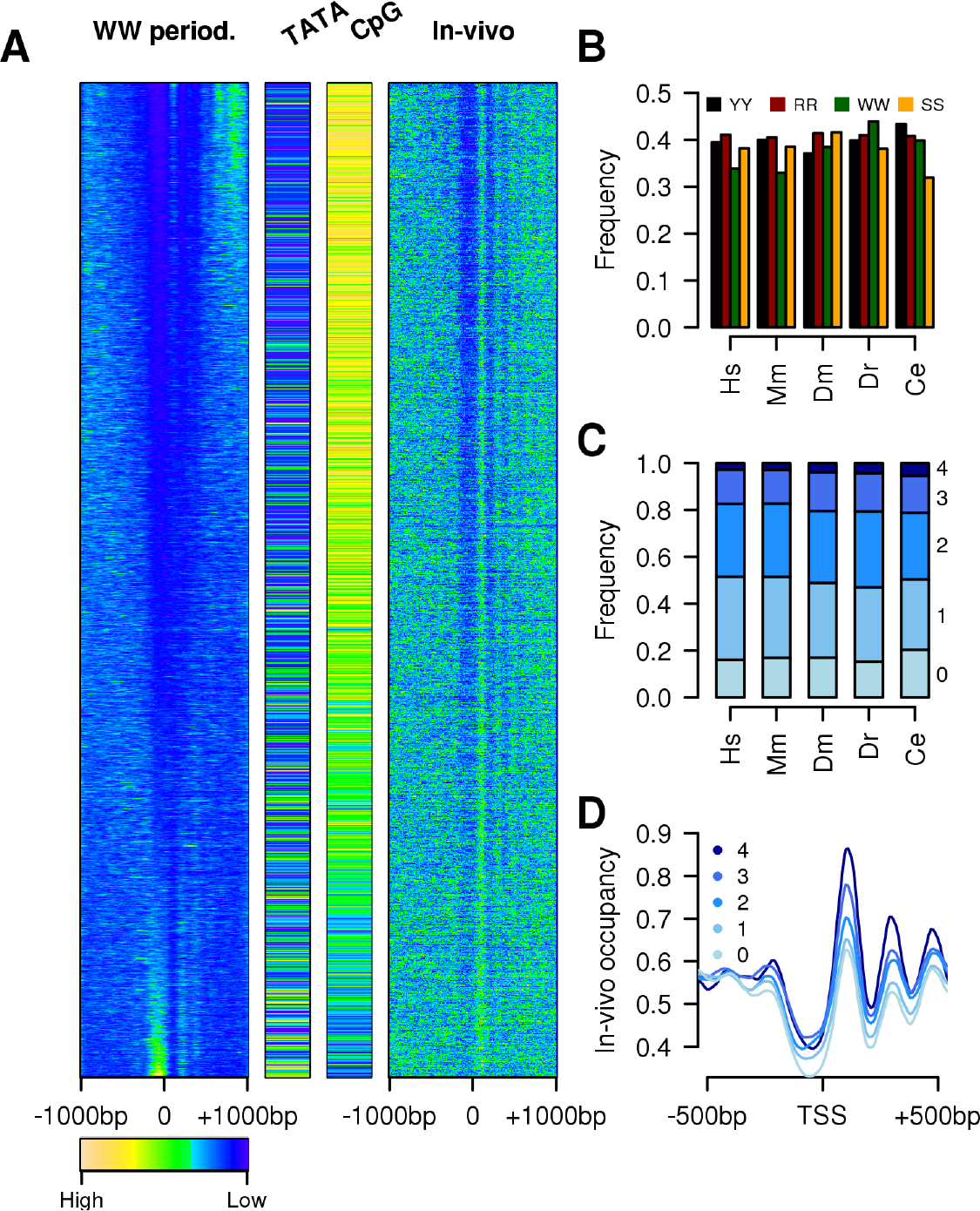
Effects of dinucleotides periodicity on chromatin. (A) Intensity of 10 bp WW periodicity calculated using a Fourier transform in a sliding window of 150 bp (10 bp shift) on a 2 kb region around all human promoters (left-hand side) compared to the *in-vivo* nucleosome occupancy profiles derived from MNase-seq reads counts in the same regions (right-hand side); Boolean TATA-box and CpG islands classifications are based on: TATA-box has to be found at position −30 to −25 bp from the TSS whereas CpG islands have to cover the TSS. Promoters were ordered according to their correlation between the intensity of the 10 bp WW periodicity and the average *in-vivo* nucleosome distribution in the region −1000 bp to 1000 bp from the TSS. Each line corresponds to the average values of 10 promoters. (B) Fraction of promoters showing low intensity of 10 bp dinucleotide frequencies in the NFR and high intensity in the N+1 region for the 4 dinucleotides tested (concordant signal). (C) Cumulative fraction of promoters (indicated with the numbers on the right) with concordant signals for the four dinucleotides. (D) Average distribution of nucleosomes around human promoters stratified by the number of their concordant signals.

### Dinucleotide periodicities have an additive effect on chromatin organisation

Fig 1A shows that a large fraction of human promoters had a WW signal that, although depleted in the NFR, did not show a clear enrichment in the N+1 region. These promoters might have had other dinucleotide signals that peaked in this region allowing for a correct nucleosome positioning. To test this hypothesis, we identified promoters with periodic signal intensity (for each dinucleotide) inthe proximal promoter region that could favour the average *in-vivo* nucleosome distribution. To do so, we compared the average 10 bp periodic signal in the NFR with that of the N+1 region and identified promoters with a higher signal downstream of the TSS (named hereafter as concordant signal). The organisms had heterogeneous number of promoters with concordant signals (Fig 1B). *H. sapiens* and *M. musculus* promoters were characterised for having the YY and RR dinucleotides as the most common and, at the same time, the WW signal was less frequent. This could have been the consequence of the presence of CpG islands that, with their high GCcontent, could affect the dinucleotide frequencies and the possibility to generate a periodic signal. WW signal was more frequent in all other organisms but only in *D. rerio* it was the most frequent. In fact, *D. melanogaster* showed that more then 40% of promoters had a concordant SS signal, whereas *C. elegans* promoters were enriched in YY signal but strongly depleted of SS signal. Nonetheless, in all organisms 80% of promoters had at least 1 concordant signal (Fig 1C) and 20% 3 or more. The presence of multiple concordant signals in the proximal promoter region was clearly reflected in chromatin organisation (Fig 1D) with more focused nucleosomes even outside the proximal-promoter area used in this analysis.

### Consensus sequences for promoters’ nucleosomes are not always similar to genomic nucleosomes

Our analyses showed that more then one dinucleotide periodic signal was often present in the N+1 region of a promoter (Fig 1). However, it was not clear how the dinucleotides were positioned compared to each other within the same sequence. The mutually exclusive WW and SS are expected to be found in counter-phases [28] as YY and RR [27]. Trifonov [45] concluded that the general DNA consensus sequence for genomic nucleosomes could be summarized with the following 2 motifs, SSRRNWWNYY or SSYYNWWNRR (note the relative position of the YY and RR in the two motifs), but little is known about the relative position of the 4 dinucleotides in the N+1 region. We addressed this using aggregate plots as in [44] where patterns of WW frequency were revealed in the N+1 region of *H. sapiens* promoters that were remarkably similar to the dinucleotide periodicities seen in MNase-seq data [28]. Using this observation, we evaluated and compared the periodic frequencies of DNA consensus sequences of the N+1 and genomic nucleosomes. To do so, promoters of the 5 organisms under study were aligned to the TSS and, using aggregated plots, the strength of a 10 bp periodic signal was evaluated in the N+1 region of all possible motifs of length 10 bp generated permuting the 4 dinucleotides and two N bases (240 motifs). A similar analysis was performed on genomic nucleosomes defined by high-resolution MNase data and aligned to the inferred center position. In *H. sapiens* (Fig 2A) there was a very high correlation between the 10 bp frequency strengths measured in DNA sequences coming from genomic nucleosomes and signal from the DNA sequences of the N+1 region with a clear separation between motifs with high signal and all the rest. Confirming the expectations from [45], motifs with strong periodicity were all characterized for having the WW dinucleotide in counter phase to the SS as well as the YY and RR and to share the same dinucleotide order: the SS dinucleotide was always followed by YY, then by WW and RR. The average intensities of this motif class around *H. sapiens* promoters showed a pattern that closely resembled *in-vivo* nucleosome maps (S5 Fig) with signal depletion in correspondence of the NFR and a peak at the N+1. Moreover, the class of motifs belonging to the first motif in Trifonov model (SS-RR-WW-YY) [45], showed very week signal in both regions. These findings indicated that in *H.sapiens*, the DNA wrapped around the histones in the N+1 region had almost identical dinucleotide periodicity patterns of the DNA found in genomic nucleosomes. *M. musculus, D. melanogaster* and *D. rerio* showed a preference for motifs belonging to the same class as *H.sapiens* (Figs 2B and S6) with a strong correlation between signals coming from genomic and promoter nucleosomes (S5 Fig). *C. elegans* was the only organism analyzed that shows a clear difference between the DNA code used on genomic nucleosome and the one used at promoters. On a genome level *C. elegans* showed no difference with the other organisms (Fig 2B left panel) with a clear preference for the motifs class SS-YY-WW-RR. *C. elegans* promoters, instead, showed strong signals also for the class SS-RR-WW-YY (Figs 2B and 2C, S5 and S7 Figs). Analysis of the average distribution of the two motif classes around *C. elegans* promoters showed signal for both (S8 Fig), suggesting the presence of two promoter groups characterized by the presence of one motif and not the other (S8 Fig). To identify them, promoters were grouped on the bases of the signal intensity for one consensus as twice as strong compared to the other. 1344 promoters had strong signal from the SS-RR-WW-YY class and 806 from the SS-YY-WW-RR. These two promoter groups did not have very different chromatin architectures with the SS-YY-WW-RR class showing only a slightly more focused N+1 and more pronounced NFR (S8 Fig) but not a difference in H3K4me3 distribution (S8 Fig).

**Fig. 2.**
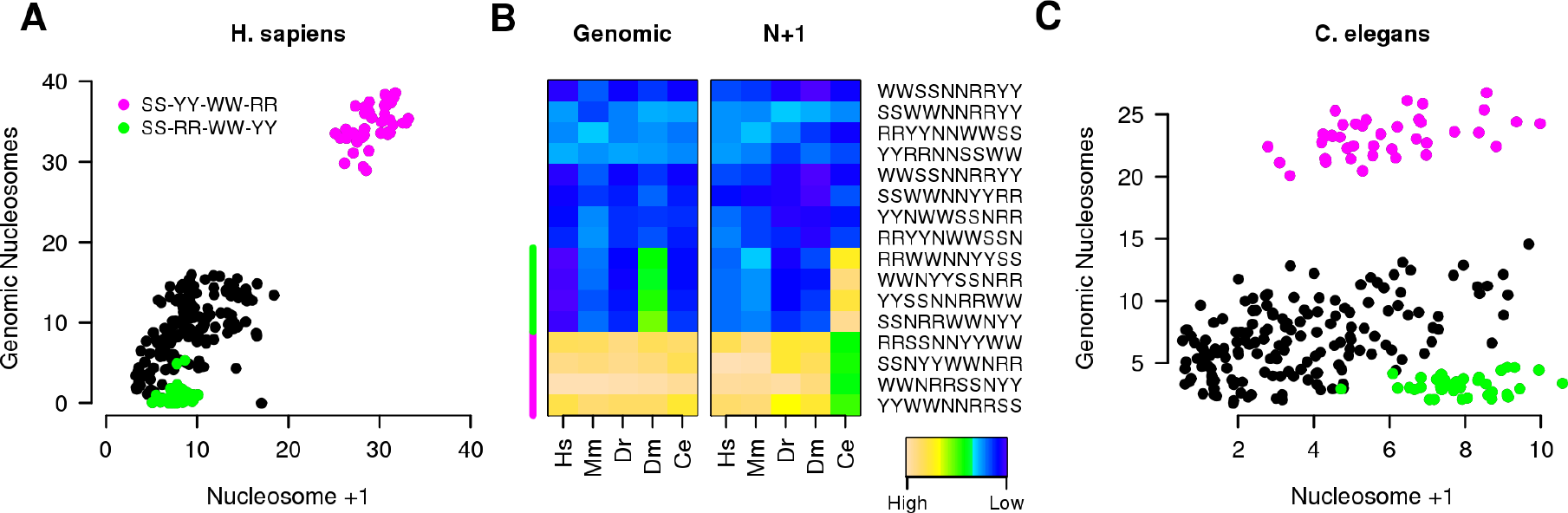
Identification of the N+1 consensus sequence. (A) Correlation between 10 bp long dinucleotide patterns composed of one copy of each SS, WW, YY and RR dinucleotides and 2 Ns evaluated on human genomic nucleosomes and the N+1 nucleosome; each dot represents the strength of the 10 bp frequency of a consensus sequence in the N+1 region or in genomic nucleosome regions (defined by MNase-seq data); green and red dots mark two classes of patterns (circular permutations of SS-RR-WW-YY and SS-YY-WW-RR, respectively) that are known to have high nucleosomes affinities [45]. (B) Comparative analysis of representative consensus sequence strength in 5 organisms on genomic nucleosomes (left panel, Genomic) and promoter nucleosomes (right panel, N+1). Each square represents the strength of a 10 bp periodic signal for a consensus sequence (rows) for an organism (columns). Hs: *H. sapiens*, Mm: *M. musculus*, Dr: *D. rerio*, Dm: *D. melanogaster* and Ce: *C. elegans.* A comparative analysis of all motifs tested can be found in S6 and S7 Figs (C) Correlation analysis as performed in A for *C. elegans* promoters and genomic nucleosomes.

### Dinucleotide periodicities in promoters correlate with Pol-II initiation patterns

The finding that promoters with a broad initiation pattern have phased dinucleotide periodicities in the N+1 region compared to focused promoters [44] that, on the other end, are enriched in TATA-box motifs [9, 17] suggests that TATA-box and chromatin conformation could have different effects on transcription initiation [12, 13]. The TATA-box can direct Pol-II to the TSS with high precision [1] whereas in its absence, chromatin organization could guide the Pol-II complex but less precisely. To analyze the quantitative effect of rotational properties of DNA on Pol-II positioning, the correlation between the strength of the dinucleotide signals in the N+1 region and the spread of Pol-II initiation were studied in grater detail. To do so, promoters were first grouped according to their TATA-box state (with and without the motif) and, for the TATA-less promoters, according to their average dispersion of Pol-II initiation around the TSS (from very focused to very broad promoters) evaluated using CAGE data and summarized with a Dispersion Index (DI, it could be considered as the standard deviation around the most likely initiation site). Then, for each group, the average strength of the four dinucleotide signals in the N+1 region was evaluated. In all organisms tested there was a strong inverse correlation between promoters DI and the average dinucleotide strength (for example for *H. sapiens*: R^2^=0.76 and p-value=0.0002) (Figs 3A and S9). Focused promoters without a TATA-box were characterized for the presence of a strong periodic signal, whereas broad promoters showed a weak periodicity. TATA-box promoters were outliers: they showed low DI values and weak periodic signals. In *D. melanogaster* another large group of promoters (5628 promoters, 1/3 of the total) was characterized for having focused initiation and weak periodicity. All these promoters had a DPE [46] and an Inr element, both of which are found at conserved distance from the TSS. Moreover in all organisms, only promoters without TATA-box (or Inr-DPE) had the signal in phase with the TSS suggesting that there was a fixed distance between the TSS and the N+1 (S10 Fig). To test if the periodic signal in the N+1 affects also the level of activity of Pol-II, the average expression of promoters was correlated with the average dinucleotide strength in the N+1 region. In this case, no correlation between the two was found (R^2^=0.18, p-value=0.21) (S11 Fig).

**Fig. 3.**
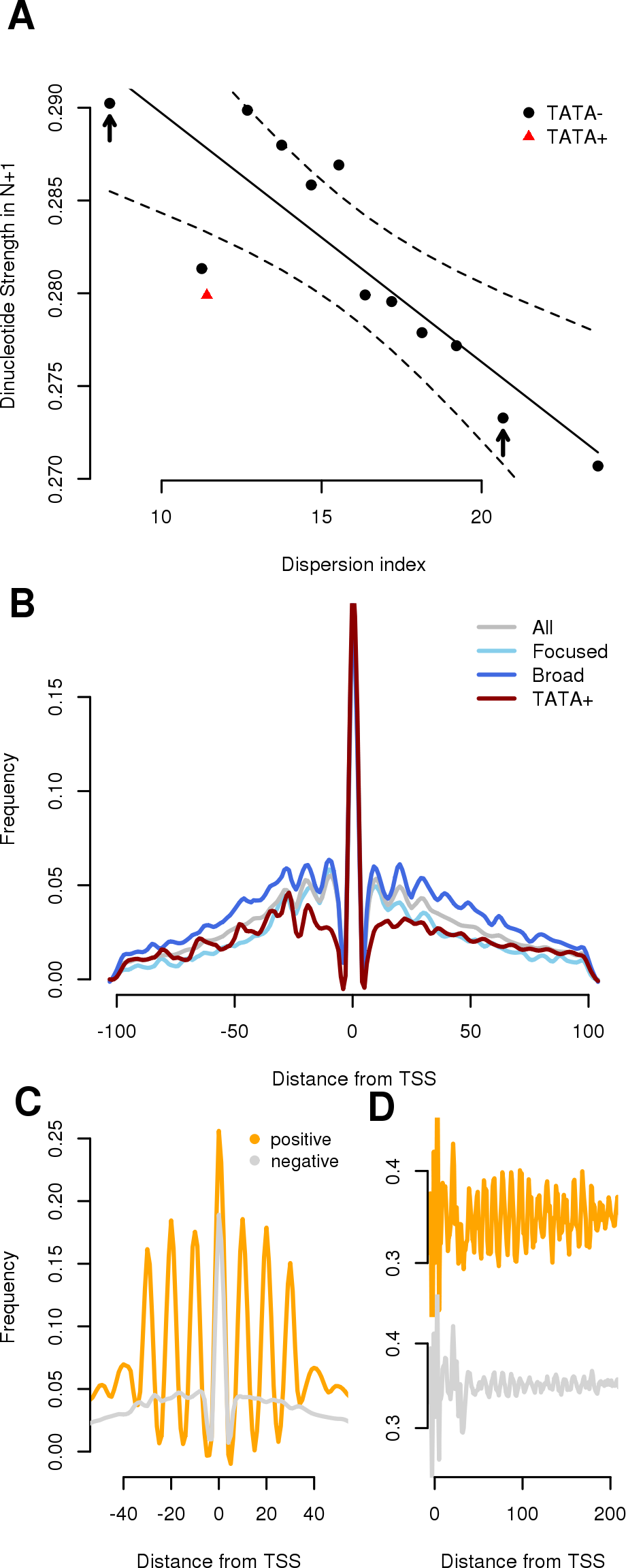
Effects of dinucleotide periodicity on Pol-II initiation. (A) Correlation between Pol-II level of dispersion of TSS initiation and strength of periodic signal in the N+1 region. Each dot corresponds to the average value of the intensity of the WW, SS, RR and YY dinucleotide of 2000 promoters ordered by increasing DI (a number similar to the TATA-box containing promoters). Solid line represents the predicted values (evaluated by a linear model) whereas dotted line the 99% confidence intervals as evaluated by the linear model. Arrows mark promoters groups with low DI (focused) and high DI (broad) used in B. (B) CAGE micro-peaks distribution around human promoters stratified for the presence or absence of TATA-box and focused or broad promoters (defined as promoter groups marked with an arrow in A). A strong 10 bp periodicity is visible only in TATA-less promoters, reflecting the presence in the N+1 region of a DNA-encoded nucleosome signal. (C) CAGE micro-peaks distribution for promoters with strong (positive) and weak (negative) periodicity in secondary initiation sites. (D) Average YY distribution for the promoter classes defined in (C).

### CPE-less promoters show 10 bp periodic initiation patterns

To further elucidate the relationship between periodic DNA signals and Pol-II, we studied the primary and secondary transcription initiation patterns in promoters. In fact, rotational nucleosome positioning due to a 10 bp periodic signal does not require the occurrence of the nucleosome center at exactly the same base: it tolerates shifting by multiples of 10 bp [26, 27]. To validate our model that the rotational setting of the +1 nucleosome influences TSS selection by Pol-II, CAGE tags were used to analyze the distribution of transcription starts at promoters. In order to detect these secondary Pol-II initiation sites, a “micro-peak” method was applied to the data that consisted in extracting positions
that corresponded to a local maximum in CAGE tag coverage within a window of 5 bp. This method emphasized the stronger initiation sites compared to a simple cut-off value and also reduced the background noise given by spurious signals (S12 Fig). Subsequently, the average distributions of secondary TSS around promoters grouped by their TATA and DI statuses were evaluated.

In *H. sapiens*, each promoter subclass showed a similar level of primary TSS activity with comparable frequencies of micro-peaks at the TSS (Fig 3B). Away from the primary TSS, two opposite Pol-II behaviors were detected. The first had a strong 10 bp periodic pattern in secondary initiation sites distribution around promoters and corresponded to TATA-less promoters regardless of their DI values with both focused and broad promoters showing strong secondary initiation patterns. The second had no clear periodic signal near the central peak and corresponded to TATA-box promoters. This subclass had also poor affinity values (Fig 3A) with the absence of a phase signal downstream the TSS (S10 Fig). The other organisms showed similar patterns of Pol-II initiation (S13 Fig) with TATA-box containing promoters the only group that did not show any periodicity in secondary initiation. In *D.* similar to TATA-box containing promoters.

The 10-bp periodic distribution of secondary initiation sites could be due to local curving of the DNA at the major initiation site or one-sided protection by components of the pre-initiation complex. To rule out this possibility and to establish a direct link between TSS phasing and the +1 nucleosome signal, we selected promoters with the strongest pattern in secondary initiation sites and studied their DNA properties in the N+1 region. Results showed that promoters with a strong periodic TSS initiation pattern (Fig 3C) also showed high phasing with the +1 nucleosome periodic signal (Fig 3D), further suggesting the presence of a direct relation between the two.

### Natural variants that map in the N+1 region alter Pol-II initiation

The strong correlation observed between DNA-encoded nucleosome positioning signals near the TSS and transcription initiation patterns (Fig 3) was an indication that the DNA sequence of promoters had a crucial role in guiding Pol-II to the initiation site via a possible N+1 interaction. To gain further evidence that there was a causative link between DNA sequence and Pol-II initiation and to identify the region that had the greatest influence, we studied the effect of natural variation on promoters’ DI. To do so, we used CAGE data from the ENCODE tier 1 cell line GM12878 (a lymphoblastoid cell line) for which the genome had been sequenced by the 1000Genome consortium [47]. Using data from this cell line, it was possible to study the effect of natural variation, such as SNPs and Indels (deletion or insertion of few bases), on Pol-II initiation expressed as variation in DI. To address this we compiled promoters’ variants for which the GM12878 was homozygous for the minor allele. In total there were 15548 SNPs mapping near promoters (2kb window around TSS) and 1849 indels. The two distributions were similar (S14 Fig), both showed low frequencies near the TSS, but were not exactly the same. SNPs minimum was centered slightly upstream the TSS whereas indels minimum downstream, in a region that coincided with the N+1.

GM12878 CAGE tags were then used to evaluate DI values for all promoters. As a reference, we used CAGE data from blood-derived cells from a different origin that should not contain the same mutations [44] and assigned them to a reference genome containing always the major allele (most likely genome). After normalization, the variation in DI of promoters with natural variants was evaluated between the two cell groups and plotted as a function of the distance of the variants to the TSS (Fig 4A). It was possible to evaluate the impact on initiation patterns made by natural variants at any given distance from the TSS. Both SNPs and indels had a measurable effect on TSS dispersion if located in the proximal promoter region. Overall, SNPs had a weaker effect on TSS dispersion, with a maximum for SNPs mapping 120 bp downstream the TSS, in the central region of the N+1 (Fig 4A). Conversely, Indels had a much stronger impact in a region that extended from the TSS until the end of the N+1 and peaked within the first half of the N+1. Interestingly, SNPs and indels mapping in the NFR did not coincide with a strong variation in DI.

**Fig. 4.**
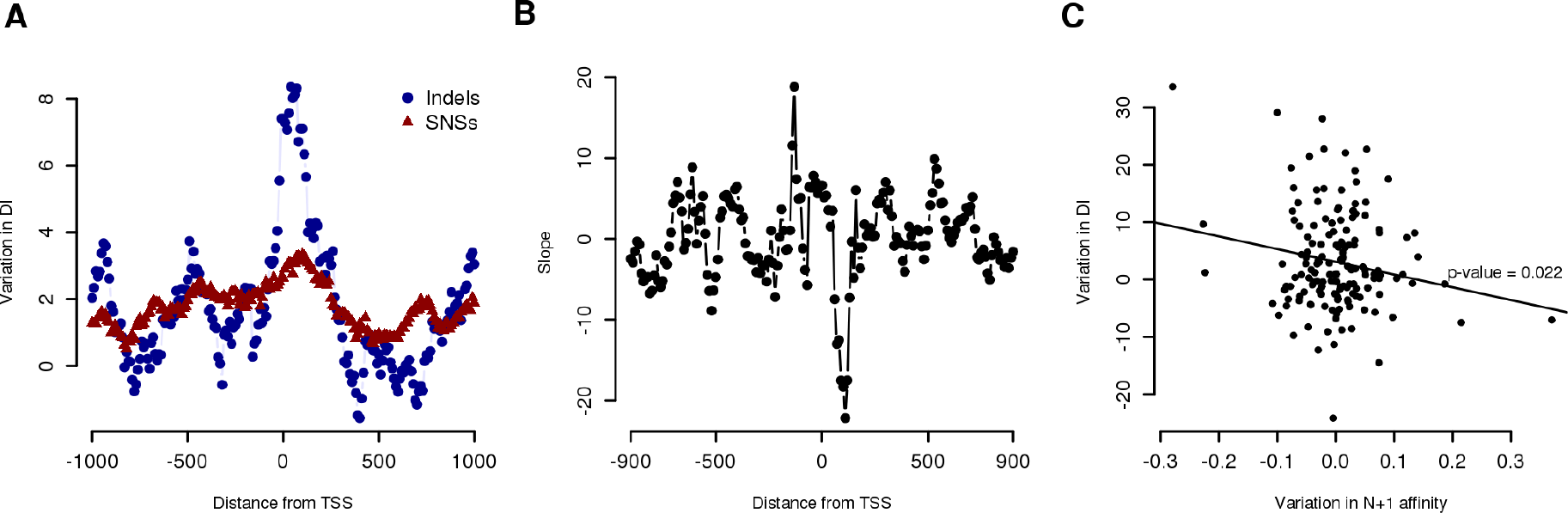
Effects of promoter natural variants on Pol-II initiation. (A) Analysis of the effect of SNPs or Indels to Pol-II initiation precision as a function of their distance from the TSS. Values on the x axis represent mid point of a sliding window of 150 bp (shift 10 bp) whereas *y* axis represent the variation in DI between GM12878 cell line and other blood derived cell lines for promoters that have natural variants in the GM12878 cell line mapping in that region. (B) 2 Kb region around human promoters were scanned for the presence of natural variants with a sliding window of 150 bp (shift 10) and analyzed for their impact on WW 10 bp frequency intensity in that region. The variation in signal intensity was then correlated to variation in DI for the corresponding promoters using a linear model. The dots represent the slope (angular coefficient) of the linear model for the region centered in that position. Negative slope values correspond to negative correlation between the variation in WW dinucleotide frequency and variation in DI (C) Effects of mutations in the N+1 region on nucleosome affinity and DI. Single promoter analysis of the effect of mutation on nucleosome affinity in the N+1 region (measured as the difference in 10 bp frequency intensity for WW dinucleotide between GM12878 and ML sequences) and its correlation with variation in DI for the same promoter (measured as difference between GM12878 and other blood related cell lines).

### Variants disrupting dinucleotide periodicity in the N+1 region tend to increase TSS dispersion

We then investigated the relationship between alterations of the nucleosomes-DNA affinity (measured as variation in dinucleotide 10 bp frequency)produced by natural variants and their effects on Pol-II initiation. To assess this, we scanned the promoter region with a sliding window of 150 bp (10 bp shift) and investigated the linear relationship between the variation in 10 bp frequency for the WW dinucleotide (produced by GM12878 natural variants that mapped in that region) and the variation in the observed DI for the corresponding promoters. The N+1 region was the only one showing a negative correlation between the variation measured in the nucleosome-DNA affinity and the variation in promoters’ DI, with a minimum centered at base +110 (p-value=0.022) (Fig 4B). On a single promoter level, natural variants that mapped in this region with disruptive effect on the nucleosome binding corresponded to promoters with increased DI compared to WT (Fig 4C). On the other end, natural variants that increased the nucleosome affinity had an effect on lowering the DI.

## Discussion

Two pathways for TSS selection by POL-II have been described in the literature. According to the conventional model the TSS position is defined by the presence of CPE [5]. However, the majority of eukaryotic promoters lack CPEs, including a TATA-box and an Inr [11]. Jiang and Pugh proposed that TSS selection in yeast might be linked to the position of the N+1 in the absence of CPEs [12]. Here, through a comparative analysis of DNA-encoded nucleosome signals in animal promoters and Pol-II initiation patterns, we report that the DNA signals underlying both mechanisms are conserved across species and, through the study of DNA natural variants, we show that the level of affinity between N+1 and DNA affects TSS selection in the absence of CPEs.

The function of sequence-intrinsic features in chromatin organization around promoters is still a matter of discussion [34]. Although studies done in yeast have shown an important role of chromatin remodeler in organizing chromatin at a genome [36] and promoter level [37], a growing body of evidence favors their active role at promoters [27, 39–41]. Here, through comparative analysis of promoters DNA sequence composition, we show that in 5 model organisms (*H. sapiens, M. musculus, D. rerio, D. melanogaster* and *C. elegans*) the position of nucleosomes at the majority of promoters is at least partly determined by DNA encoded signals, with some remarkably species-specific differences. Promoters of all organisms show a 10 bp periodic signal for the four dinucleotides tested (WW, SS, YY and RR). *H.sapiens* is the only organism showing also a strong signal for YY and RR dinucleotides for a periodicity of 8 bp, that is probably the consequence of the presence of specific CT rich microsatellite sequences in human promoters [48] (S1 Fig). As expected, the dinucleotide that shows the highest correlation with *in-vivo* nucleosome maps is WW (Fig 1A). Regardless of this, multiple periodic signals reinforce each other in organizing chromatin around promoters (Fig 1D), suggesting an additive effect of the affinity of the four dinucleotides to histones. When we study the spatial relationships between the four dinucleotides within a promoter sequence we find the same consensus as in genomic nucleosomes (SS-YY-WW-RR) for all organisms tested with the exception of *C. elegans.* Interestingly, on a genome level the DNA that is wrapped around *C. elegans* nucleosomes has the same consensus sequence as all other organisms but at promoter level we find that there are two distinct group of promoters characterized for having the SS-YY-WW-RR or SS-RR-WW-YY consensus. This finding is intriguing since the difference in the two sequences is not purely semantic, but has been predicted to alter the affinities to histones [45]. Although SS-RR-WW-YY has been predicted to have the higher affinity to nucleosomes allowing for perfect bendability of the DNA around the histone octamer [49], our analysis show that *C. elegans* promoters with this sequence in the N+1 region do not have any difference in chromatin conformation compared to promoters with the other consensus. The reason for this unexpected observation is unknown and need further investigation.

The identification of promoters by the transcription machinery is a process that is guided by the general transcription factor TFIID [50], a multi-subunit protein that is not only able to interact with the TATA-box or the DPE element [5] but also with chromatin [51–53] via the TAF3 subunit, suggesting the presence of a motif-independent TFIID recruitment at promoters that rely on the N+1 [54]. In agreement with this hypothesis, TATA-box mutation studies have shown a direct effect on Pol-II initiation both in term of TSS position and level of promoter activity [19, 55]. On the other end, no study, to our knowledge, has investigated the effect that nucleosome-DNA affinity in the N+1 region has on TSS selection. Correlation analysis shows that in all organisms promoters without CPEs have the predicted level of nucleosome-DNA affinity anti-correlated with TSS initiation patterns (Figs 3A and S9). Broad promoters generally have lower DNA-encoded nucleosome affinity. Conversely, narrow promoters, often presented as a homogeneous class in the literature, vary greatly in this respect, with only the CPE-less subset (TATA-less and Inr-DPE-less in *D. melanogaster*) showing strong affinity in the N+1 region. Moreover, the 10 bp periodicity seen in Pol-II initiation in all promoters, focused and broad, that lack CPEs (Figs 3B and S13) is another indication of a direct interaction between Pol-II and the N+1 in the absence of other DNA signals. In fact, a model of Pol-II initiation that relies on the interaction with the N+1, which in turn is rotationally positioned and able to tolerate shifting by multiples of 10 bp [26, 27], would allow Pol-II to start transcription at 10 bp intervals. Furthermore, the study of DNA natural variants in *H.sapiens* have shown that the region with grater influence on TSS selection is the N+1 (Fig 4A) and that there is a negative correlation between variation in nucleosome affinity and Pol-II initiation (Fig 4 panels B and C). That is, the presence of a variant in the N+1 region that decreases the nucleosome-DNA affinity results in an increase in TSS dispersion and vice-versa. These results strongly support the model of a motif-independent TFIID recruitment mediated by N+1—TAF2 interaction [54]. We can speculate that, in the absence of the TATA-box or Inr-DPE, the relative stability of the histones-DNA complex in the N+1 region could be transferred to the PIC via interaction with TFIID leading to a more or less focused transcription initiation by Pol-II. On the other end, in all organisms studied, CPEs containing promoters are outliers compared to non-CPE promoters: they are focused but have weak nucleosome affinity and do not show any TSS periodicity. In this class of promoters the initiation site appears to be specified solely by the presence of the CPE [8, 10].

## Methods

The study is based on experimental evidence present in public datasets. All arithmetic computations were done in R and the corresponding code is presented in the Data Reproduction Guide provided as supplementary material. This document follows high standards of reproducible research; it is a step-by-step guide to precisely reproduce all results presented in this paper and to generate all the figures.

### Data Sets

The promoter sets and the corresponding dominant TSS positions were taken from EPDnew [11]: version 2 for *H.sapiens* and *D. melanogaster*, version 1 for all other species. Pol-II initiation patterns were based on CAGE or GRO-Cap data from the following sources: *H. sapiens*: ENCODE data, GEO ID GSE34448 [56], FANTOM5 [44]; *M. musculus*: FANTOM5 [44]; *D. rerio* SRA ID SRA055273 [57]; *D. melanogaster* SRA ID SRP001602; *C. elegans* GRO-cap data GSE43087 [58].

Nucleosome maps are from paired-end MNase-seq data or alternatively from single-end MNase-seq data. *H. sapiens*: paired-end MNase-seq data for the lymoblastoid cell line GM18507, SRA ID SRP012024, GEO ID GSM907783 [28], *M. musculus*: single-end MNase data from HAFTL cell line, GEO-ID GSM1293995 [59]; *D. rerio*: single-end MNase-seq data from embryos in dome stage, GEO ID GSM1081554 [60]; *D. melanogaster*: paired-end MNase-seq data, GEO ID GSM1293957 [61]; *C. elegans*: paired-end MNase data from adults, SRA ID SRP000191 [62].

### Position weight matrices for CPEs and CpG island annotation

Promoter lists were stratified based on the presence or absence of core promoter elements using the TATA-box and Inr position weight matrices (PWMs) from [6]. Promoter sequences were scanned with these PWMs using the cut-off values suggested in the original paper. Promoters were classified as TATA+ if a TATA-box was present at position-29±3 relative to the TSS, while as Inr+ if this motif occurred exactly at the TSS. The *D. melanogaster* Inr-DPE matrix is posted at http://epd.vital-it.ch/promoter_elements/init-dpe.php, including the recommended cut-off values.

CGI coordinates for human and mouse were downloaded from the UCSC genome browser [63]. Promoters with a CGI that spans the TSS (starting before and ending after the TSS) were attributed to the CGI+ class.

### Evaluation of periodicity score around promoters

Promoter sequences from position −1074 to position 1075 relative to the TSS were extracted from the corresponding genome assembly (*H. sapiens*: hg19; *M. musculus*: mm9; *D. rerio*: danRer7; *D. melanogaster*: dm3; *C. elegans*: ce6) and scanned for the presence of four dinucleotide types (identified by IUPAC codes): WW (W=A or T), SS (S=C or G), RR (R=A or G) and YY (Y=C or T). The resulting binary sequences were individually scanned in a sliding window of 150 bp, shifted by 10 bp at a time. A Fourier transform was applied to each window in order to extract the power spectrum. From the resulting spectrum, the value corresponding to a frequency of 0.097 (corresponding to a period of 10.3 bp) was extracted. This value was directly used as a periodicity score.

### Identification of genomic nucleosomes

For paired-end samples, nucleosome positions were restricted to paired-reads that formed fragments of exactly 147 bp as previously reported in [28]. In a similar way, to reproduce analogous results on single-end samples, reads were selected if they had another read mapped on the opposite strand 147 bp downstream. For both single-and paired-end samples, multiple fragments that mapped to the same location were considered only once. For both paired-and single-end samples, the midpoints of the fragments were used as the inferred nucleosome position.

### Evaluation of consensus motifs scores for nucleosome +1 and genomic nucleosomes

Consensus motifs were generated by permuting the 4 dinucleotide (WW, SS, YY, RR) and two Ns. Sequences starting with an N were discarded resulting in a total of 240 sequences. These consensus motifs were then mapped to promoters and MNase-seq enriched regions.

For the analysis of nucleosome +1, the region from position-99 to 300 relative to the TSS of the corresponding genome assembly was used for mapping each consensus motif allowing a maximum of 3 mis-matches. Then, the average occurrence frequency for each motif was evaluated from base +50 to +200 relative to the TSS and a Fourier transform was applied in order to identify the intensity of the frequency of 0.097 (corresponding to a period of 10.3 bp). This value was then stored as the motifs’ score for the nucleosome +1 and the procedure was repeated for all consensus motifs. For the genomic nucleosomes a similar analysis was performed. In order to speed-up the analysis, 80.000 inferred positions were randomly selected from each sample. Subsequently, each consensus motif was mapped around the inferred nucleosome position and the average occurrence frequency was calculated from position-75 to +75 relative to it. A Fourier transform was then applied as before and the value for a period of 10.3 bp was used as the motif score in genomic nucleosomes.

### Periodicity analysis of Pol-II initiation patterns

CAGE data from different samples belonging to the same species were first merged into one file. TSS profiles were then extracted for promoter regions extending from-103 to +104 relative to the dominant TSS using the ChIP-Extract tool from the ChIP-Seq web server [64]. The resulting integer arrays were then converted into binary “micro-peak” arrays. Briefly, a micro-peak corresponds to a 5bp window with a minimal number of 100 tags. The position of the micro-peak is then assigned to the position with the highest number of tags within the corresponding window. Each micro-peak was then given a maximum value of 1 tag. The cumulative frequency of micro-peaks was then determined at singlebase resolution within a 200bp region around the TSS.

To identify promoters with a strong 10 bp periodicity in micro-peaks signals, promoters were ranked according to the covariance between their micro-peaks distribution and a cosine function of period 10 bp. Promoters with weak micro-peak signal (with low covariance values) were selected for having a cumulative covariance equal to 0.

### Nucleosome Distribution Around Promoters

Nucleosome distributions for promoter subsets were computed from nucleosome mapping data using the ChIP-Cor program from the ChIP-Seq web server [64]. MNase-or ChIP-seq tags were centered by 70 bp to account for the estimated fragment size of about 140 bp (centering parameter of the ChIP-Seq server). Multiple tags mapping to the same genomic location were removed from the analysis (parameter “Count cut-off” set to 1) and tag frequencies were calculated in a 10 bp sliding window.

### Evaluation of Dispersion Index (DI)

The spread of CAGE tags in a window of 100 bp around the TSS was expressed as a Dispersion Index (DI) using the following formula:

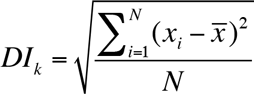

Where *N* is the total number of tag starts in the window around promoter *k*, and *x_i_*, is the mapped position of the 5’ end of tag *i.* For each species, DI values were calculated for each promoter using CAGE data from individual samples. A DI was calculated only if more then 5 tags mapped in the selected region. The sample–specific DI were then averaged to obtain a final unique and robust DI value for each promoter.

### Analysis of genomic variants in GM12878 cell line and generation ofa Most Likely (ML) genome

VCF files of Indels (version 2010_07) and SNPs (version 2010_03) for the GM12878 cell line were downloaded from the 1000Genomes ftp web server. All homozygous variants were extracted from these files and used to generate a GM12878 genome. On the other end the frequencies of these variants were evaluated using the allele frequency calculated by the final version of the 1000Genome project (phase 3, 20130502). For each variant, the most frequent allele was stored and used to generate the Most Likely genome that was then used as reference. The final list of SNPs and Indels for GM12878 cell line was restricted to the variants that differ compared to the ML genome.

## Acknowledgments

We thank Debora Gasperini for critical comments on the manuscript.

## Author Contributions

Conceived and designed the analyses: RD and PB; Performed the analysis: RD; wrote the software: RD and AG; wrote the paper: RD and PB.

## Supporting Information

**S1 Fig.**
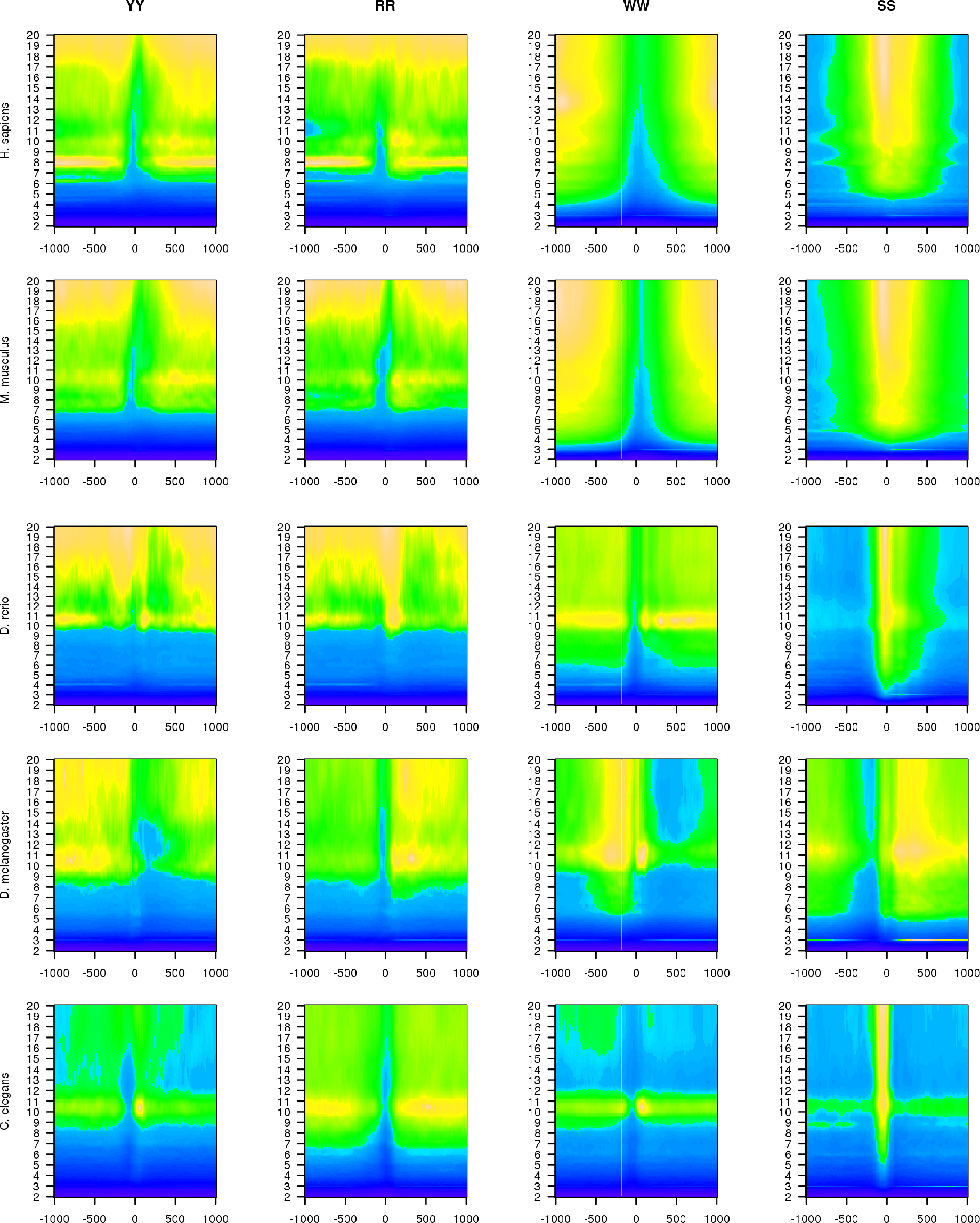
Periodic signal intensity around animal promoters. 2 kb region around promoters were scanned with a sliding window of 150 bp and 10 bp shift for the intensity of dinucleotide frequencies of period 2 to 20 bp. For each region, the average frequency intensities across all promoters were plotted against the distance of the region to the TSS. All organisms have a peak of signal intensity in correspondence to a period of 10-11 bp in agreement with the notion that this frequency helps the DNA wrapping around the histone octamer.

**S2 Fig.**
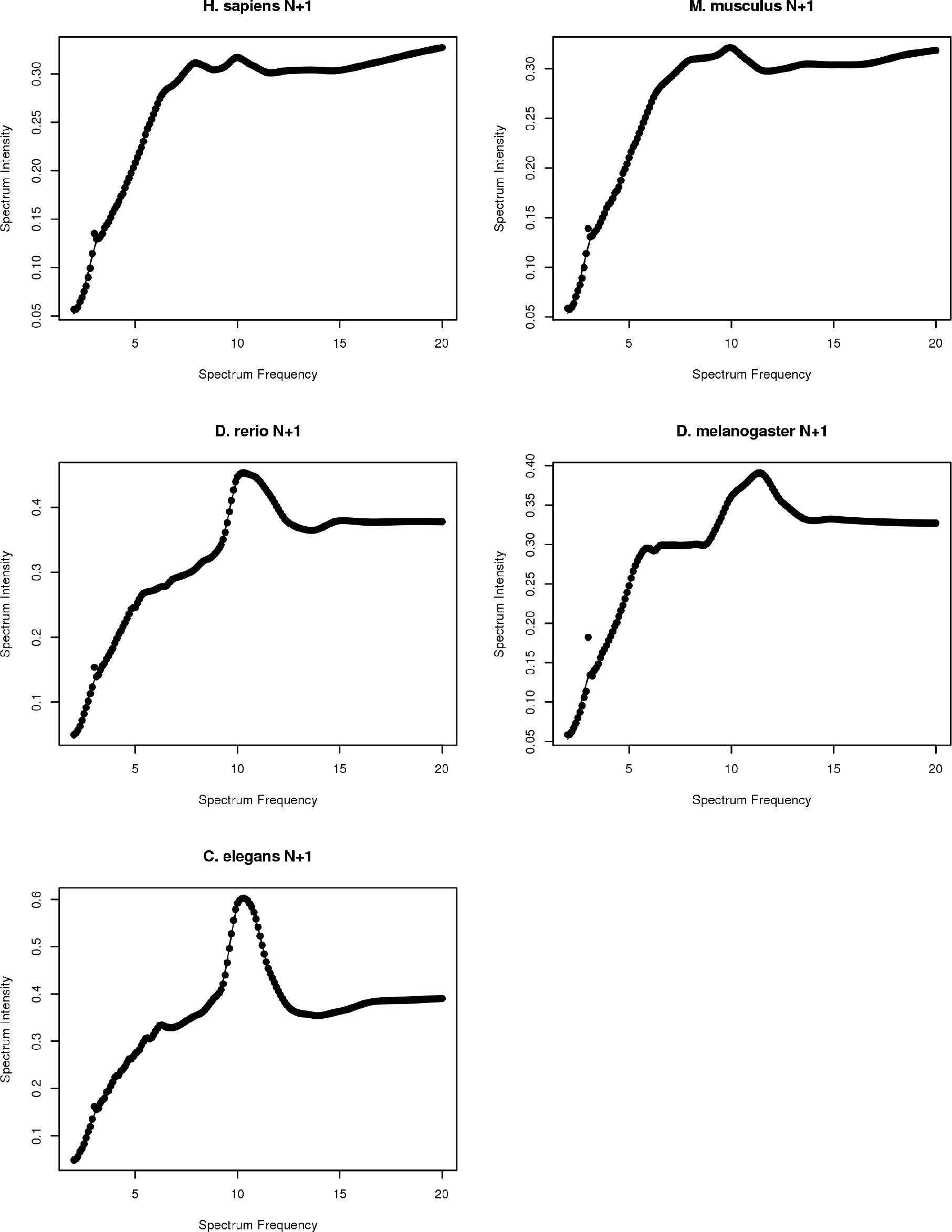
Spectrum intensities in the N+1 region. Average spectrum intensities for selected dinucleotides evaluated at position +120 bp from the TSS. Each organism shows a peak in correspondence of 10-11 bp frequency. Dinucleotide selected: *H.sapiens* and *M. musculus*: RR; *D. rerio, D. melanogaster* and *C. elegans*:WW.

**S3 Fig.**
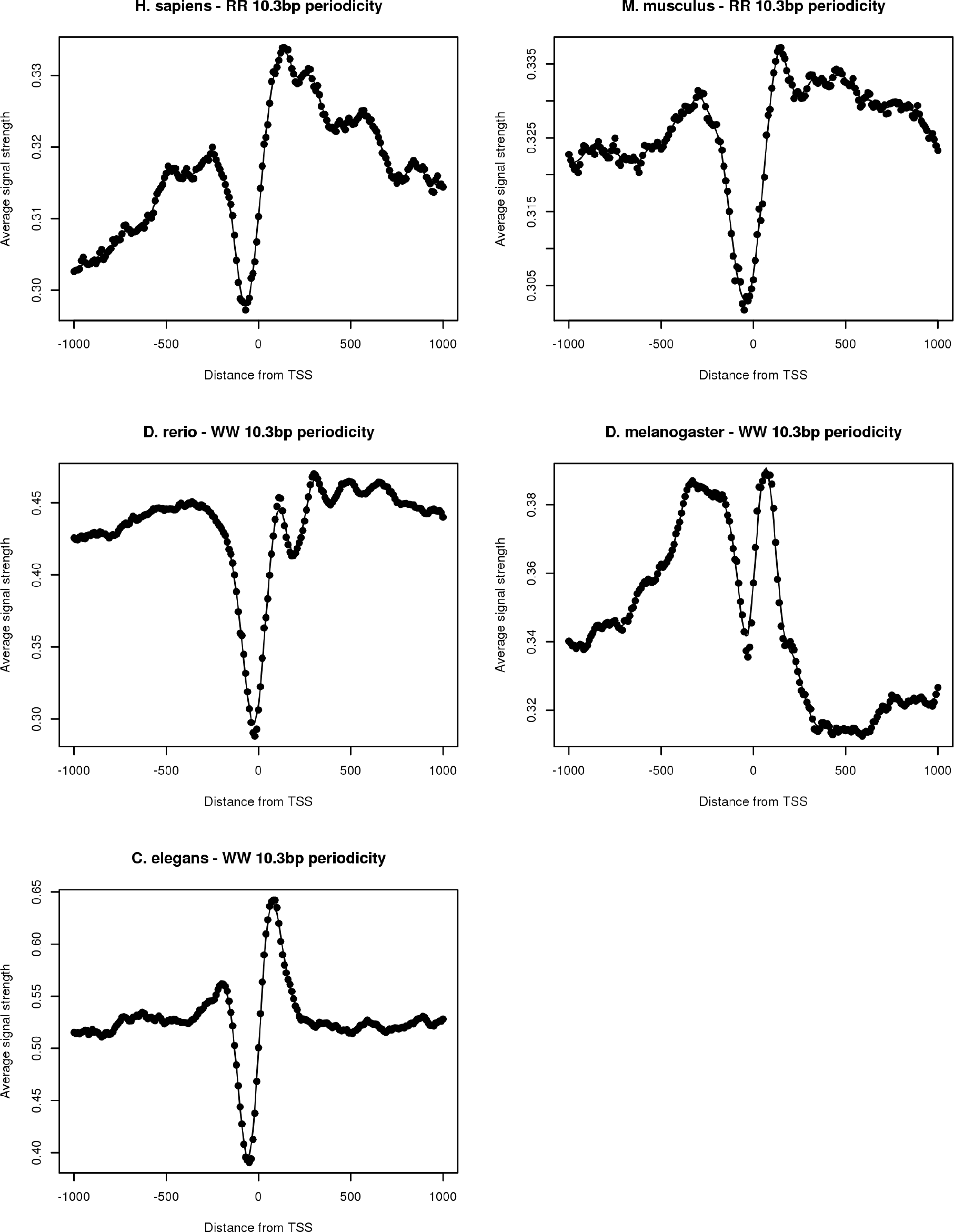
Dinucleotide 10 bp frequency around promoters. The average intensity of 10 bp frequency for selected dinucleotides in a 2 Kb region around animal promoters. All organisms show signal depletion immediately upstream the TSS followed by a peak downstream.

**S4 Fig.**
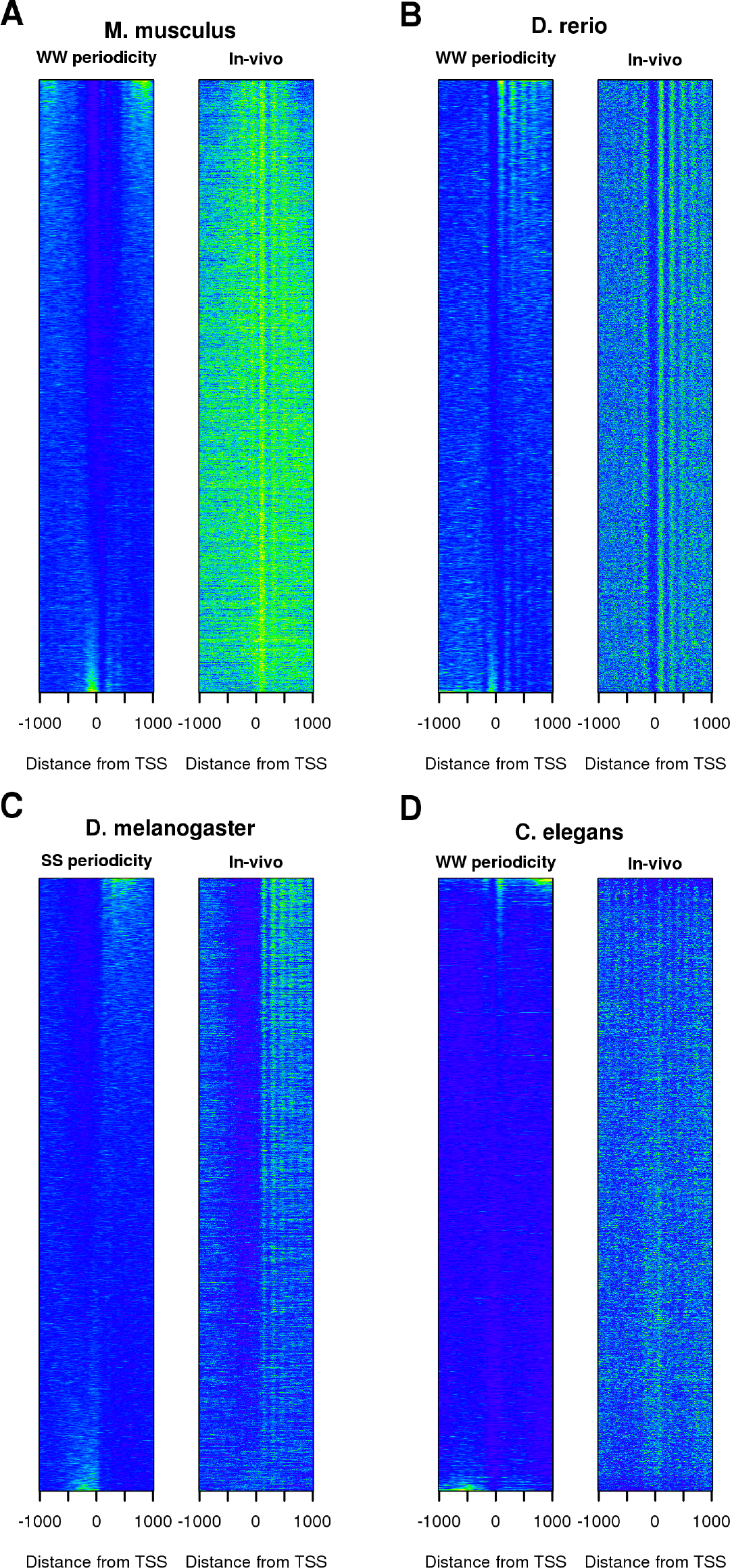
Effects of dinucleotides periodicity on chromatin. Intensity of a 10 bp dinucleotide periodicity calculated using a Fourier transform ina sliding window of 150 bp on a 2 kb region around animal promoters compared to the *in-vivo* nucleosome occupancy profiles derived from MNase-seq reads counts in the same region. Promoters were ordered according to their correlation between the intensity of the 10 bp dinucleotide signal and the average *in-vitro* nucleosome distribution in the same region.

**S5 Fig.**
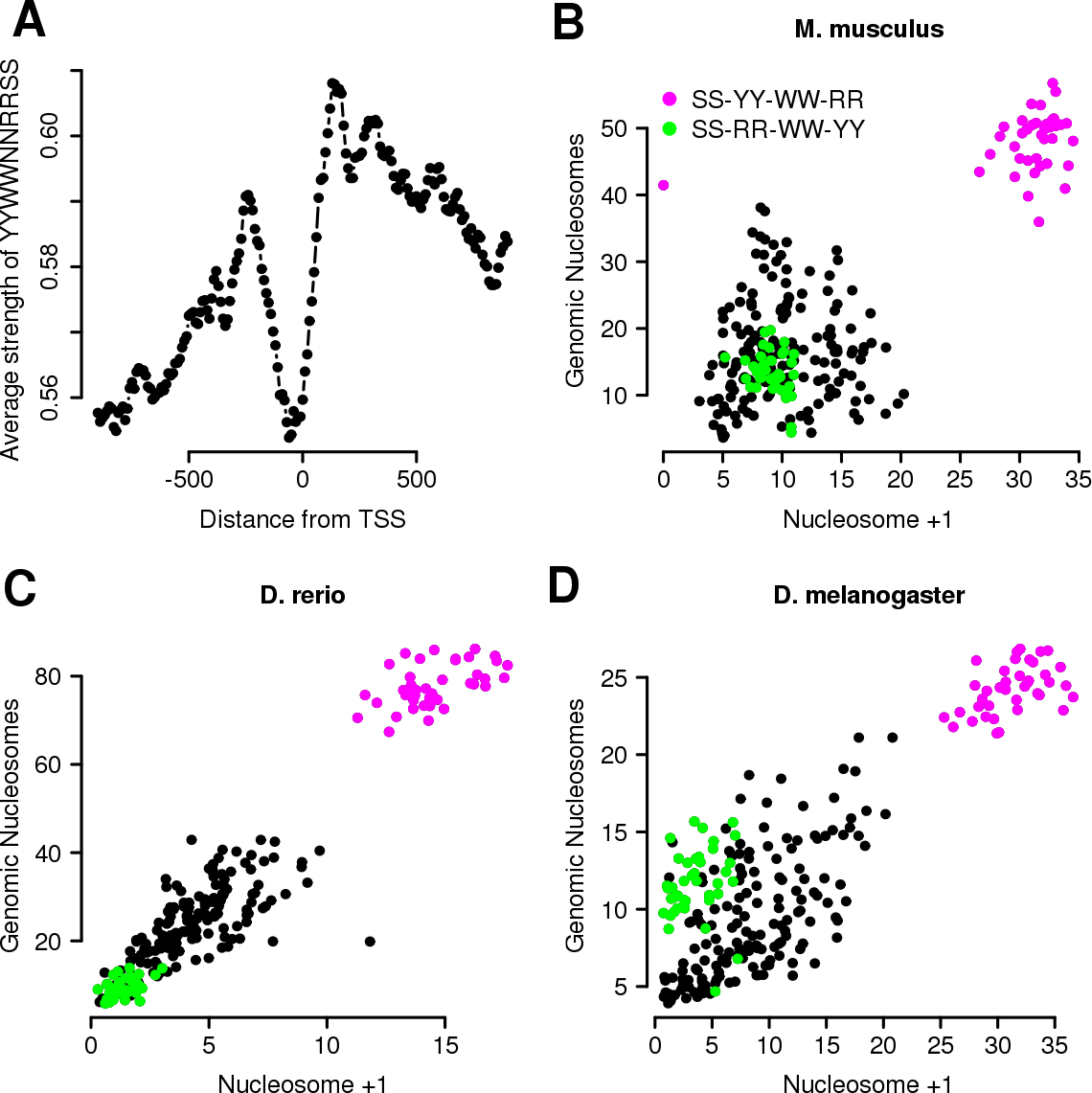
Identification of the N+1 consensus sequence. (A) Average 10 bp frequency intensity of the consensus sequence YYWWNNRRSS (3 mismatches allowed) around *H.sapiens* promoters. (B) Correlation between 10 bp long dinucleotide patterns composed of one copy of each SS, WW, YY and RR dinucleotides and 2 Ns evaluated on *M. musculus* genomic nucleosomes and the N+1 nucleosome; each dot represents the strength of the 10 bp frequency of a consensus sequence in the N+1 region or in genomic nucleosome regions (defined by MNase-seq data); green and red dots mark two classes of patterns (circular permutations of SS-RR-WW-YY and SS-YY-WW-RR, respectively) that are known to have high nucleosomes affinities. (C) As B) but for *D. rerio* promoters and genomic nucleosomes. (D) As for (B) but for *D. melanogaster* promoters and genomic nucleosomes

**S6 Fig.**
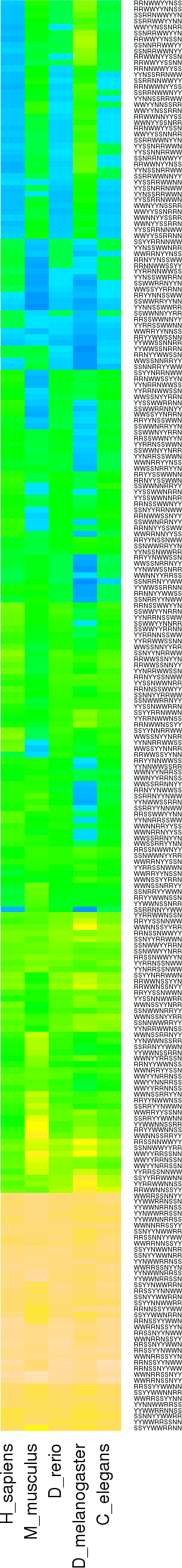
Intensity of consensus sequences in genomic nucleosomes. 10 bp frequency intensity for the 240 randomly generated sequences in MNase-seq defined genomic nucleosomes for the 5 organisms under study.

**S7 Fig.**
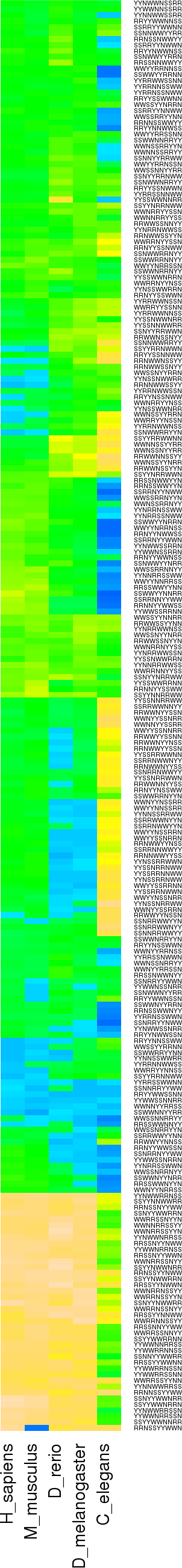
Intensity of consensus sequences in promoter nucleosomes. 10 bp frequency intensity for the 240 randomly generated sequences in the region +50 to +200 from the TSS of the organisms tested.

**S8 Fig.**
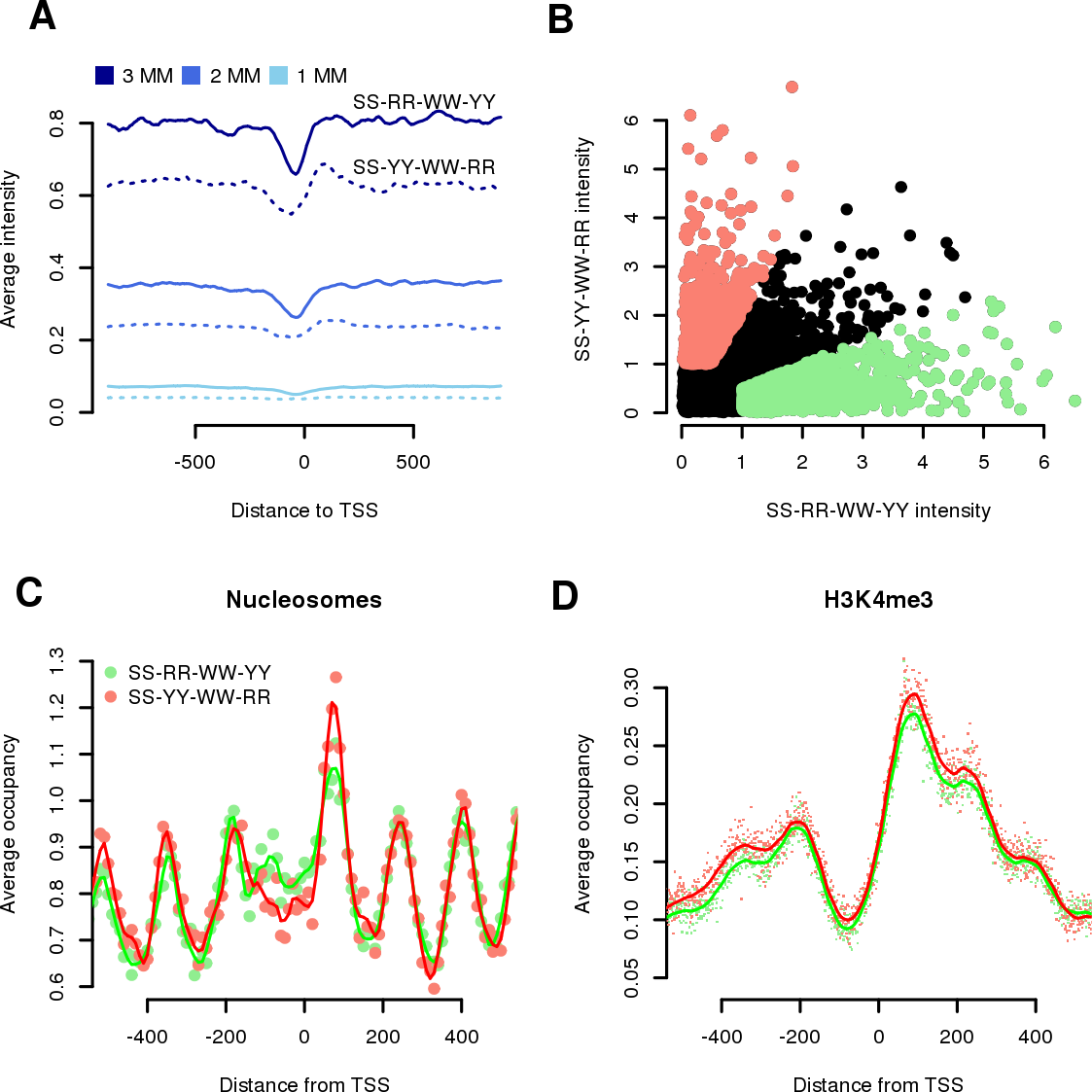
Analysis of the two consensus sequences in *C. elegans* promoters. (A) Average 10 bp frequency intensities of two consensus classes (represented by the RRWWNNYYSS and YYWWNNRRSS consensuses for the SS-RR-WW-YY and SS-YY-WW-RR classes) around *C. elegans* promoters. Different shades of blue represent the total number of mismatches allowed in the mapping. (B) Intensity of the 10 bp frequency of the two consensuses on the N+1 region for each *C. elegans* promoters. Red and green dots highlight promoters characterized for a strong signal of the YYWWNNRRSS consensus (double the signal) compared to the RRWWNNYYSS consensus respectively. (C) Nucleosome distribution around promoters characterized for a strong signal in one consensus as defined by (B). Each dot represents the average tag count in a widow of 10 bp. Continuous lines are the local polynomial regression fit. (D) Same as (C) but with H3K4me3 and window of 1 bp.

**S9 Fig.**
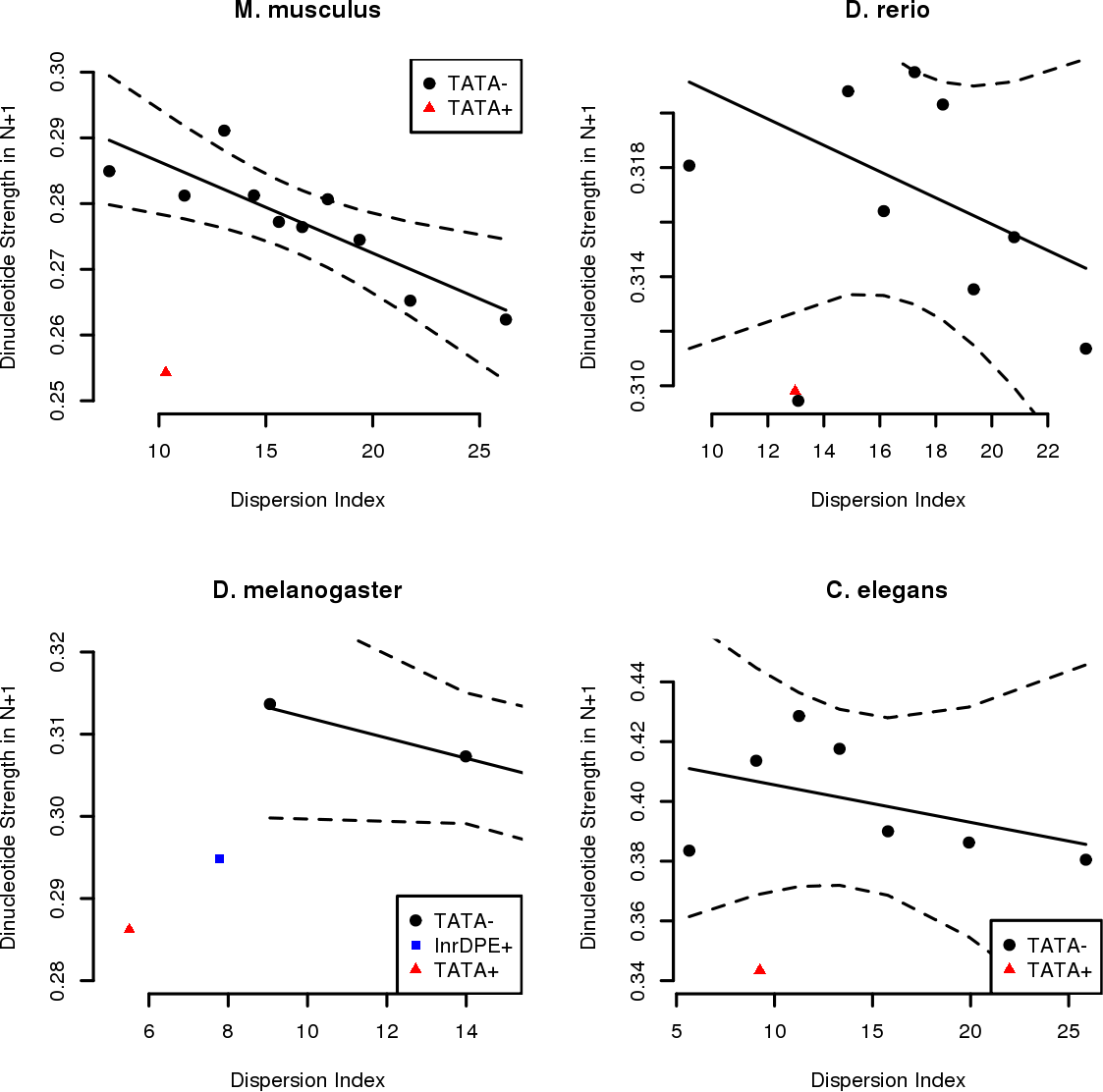
Correlation between Pol-II level of dispersion and the strength of periodic signal in the N+1 region. For each organism promoters were grouped according to their CPE status (TATA-box and Inr-DPE presence in the expected position). CPE-less promoters were also grouped according to their DI value in groups of 2000 promotes (a similar number as the promoters with the TATA-box). Each dot represents the average value of the intensity of the WW, SS, RR and YY dinucleotide for groups with similar DI and CPE status. Solid line represents the predicted values (evaluated by a linear model) whereas dotted line the 99% confidence intervals as evaluated by the linear model.

**S10 Fig.**
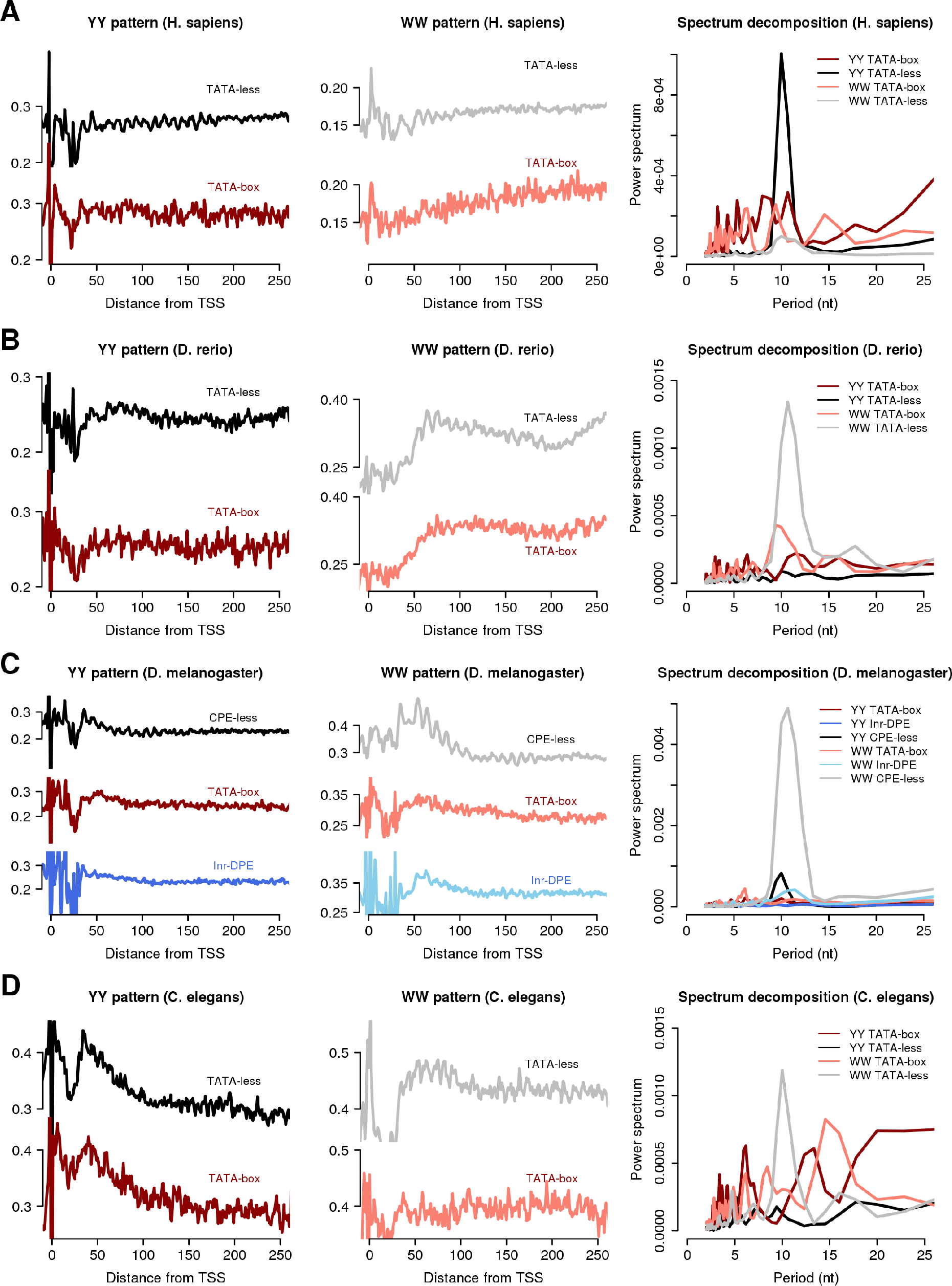
Phase-relationship of TSS and dinucleotide frequencies in the nucleosome +1 region stratified for promoters CPE statuses. (A) YY dinucleotide frequencies for *H.sapiens* promoters with TATA-box (TATA-box) and without (TATA-less) (left panel); WW dinucleotide frequencies for the same promoters groups (central panel); and spectrum decomposition in the region +50 to +200 (right panel) for the 4 signals. (B) Same as A but for *D. rerio* promoters. (C) Similar to A but with *D. melanogaster* promoters stratified for the presence of the TATA-box (TATA-box), the absence of the TATA-box but the presence of Inr-DPE motif (Inr-DPE) and for the absence of both (CPE-less). (D) Same as (A) but with *C. elegans* promoters.

**S11 Fig.**
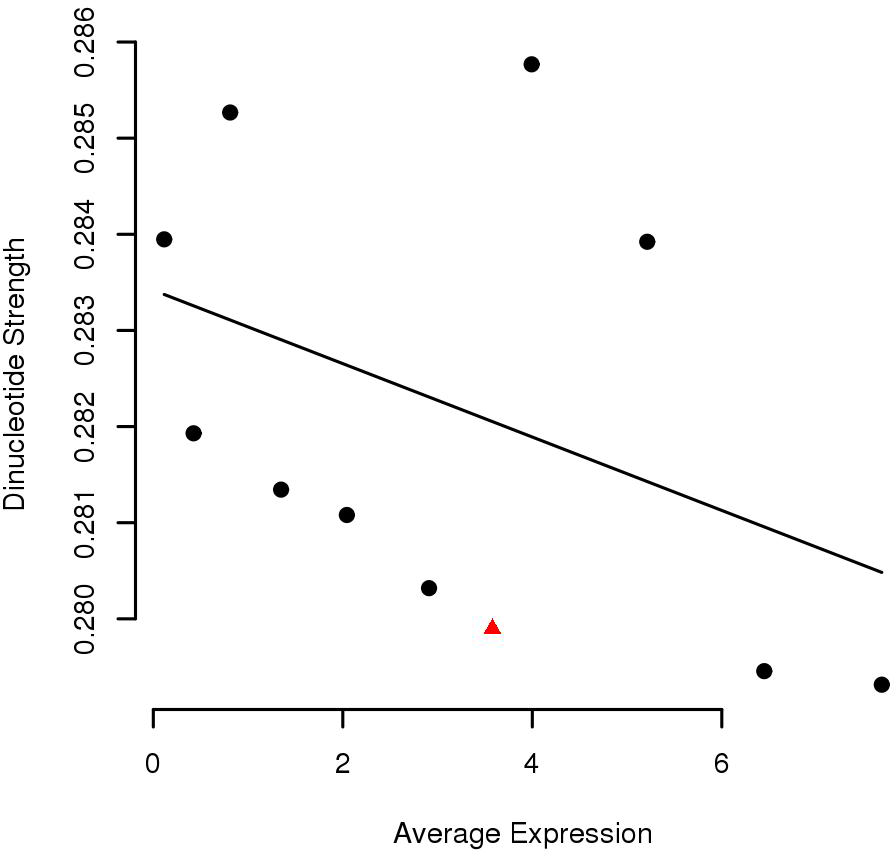
Correlation analysis between average promoter expression and strength of periodic signal in the N+1 region. *H.sapiens* promoters were grouped following the same rules as in S9 Fig but using the average promoter expression instead of Dispersion Index. In this case, no correlation is seen between expression and average dinucleotide 10 bp frequency strength in the N+1 region.

**S12 Fig.**
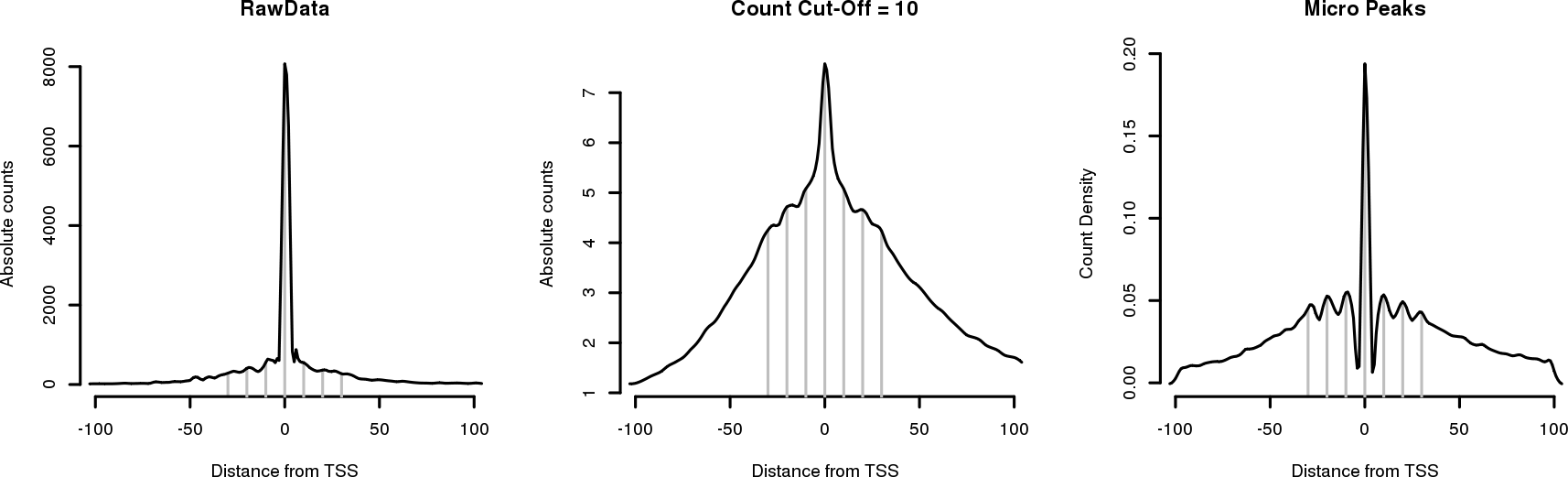
Effect of different normalization methods on CAGE distribution around *H.sapiens* promoters. Vertical grey lines are separated by 10 bp from position-30 to 30 relative to the dominant TSS. Left panel: raw global CAGE distribution, periodic initiation is not clearly visible. Middle panel: CAGE distribution after a 10 tags count cut-off was applied to each position around each promoter, a 10 bp periodicity is starting to emerge from the data. Right panel: micro peak distribution, 10 bp periodic distribution is evident.

**S13 Fig.**
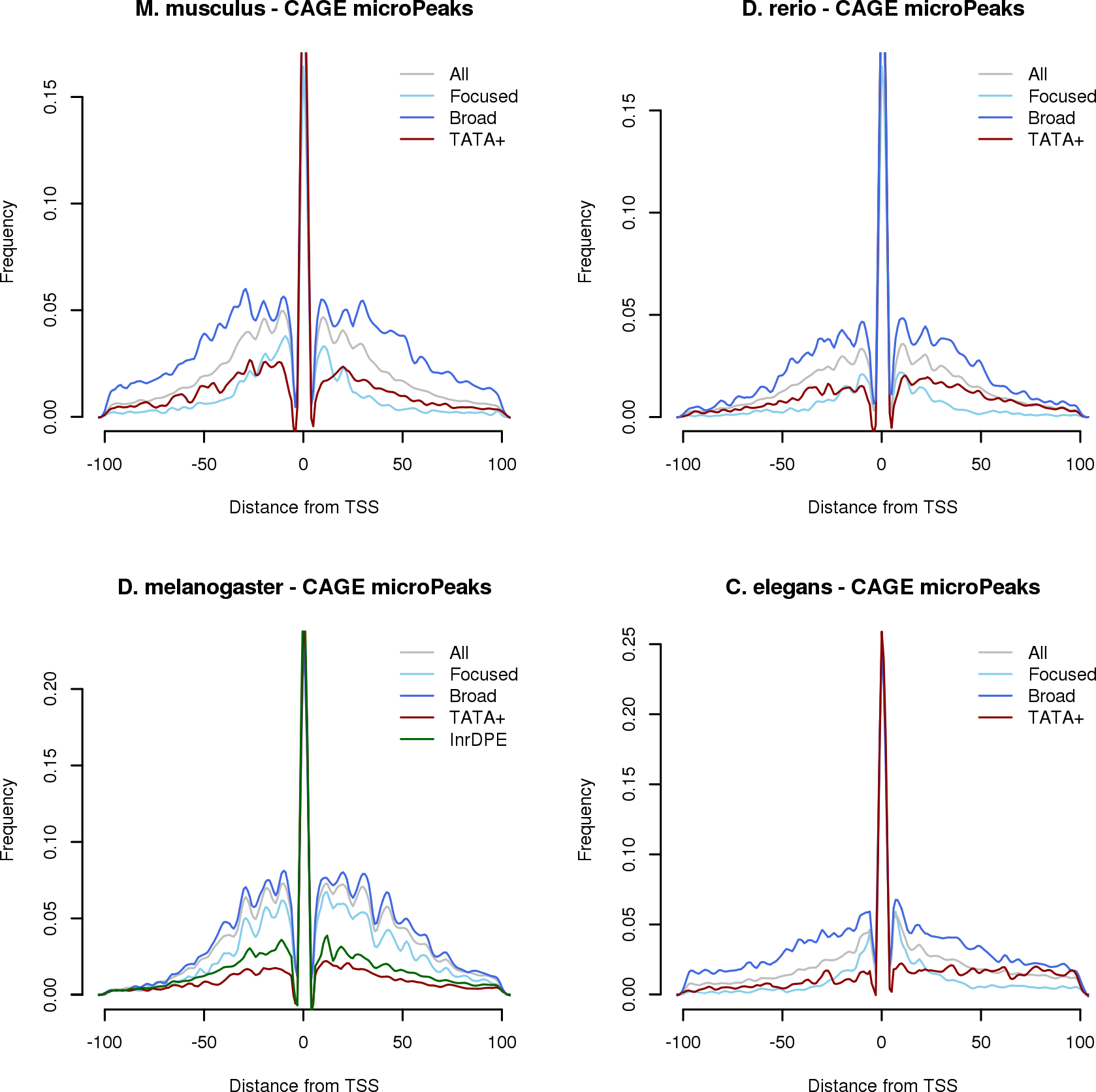
CAGE micro-peaks distribution around promoters stratified by their CPE status and Pol-II initiation pattern. A strong 10 bp periodicity in Pol-II initiation is visible only in CPE-less promoters, reflecting the presence in the N+1 region of a DNA-encoded nucleosome signal.

**S14 Fig.**
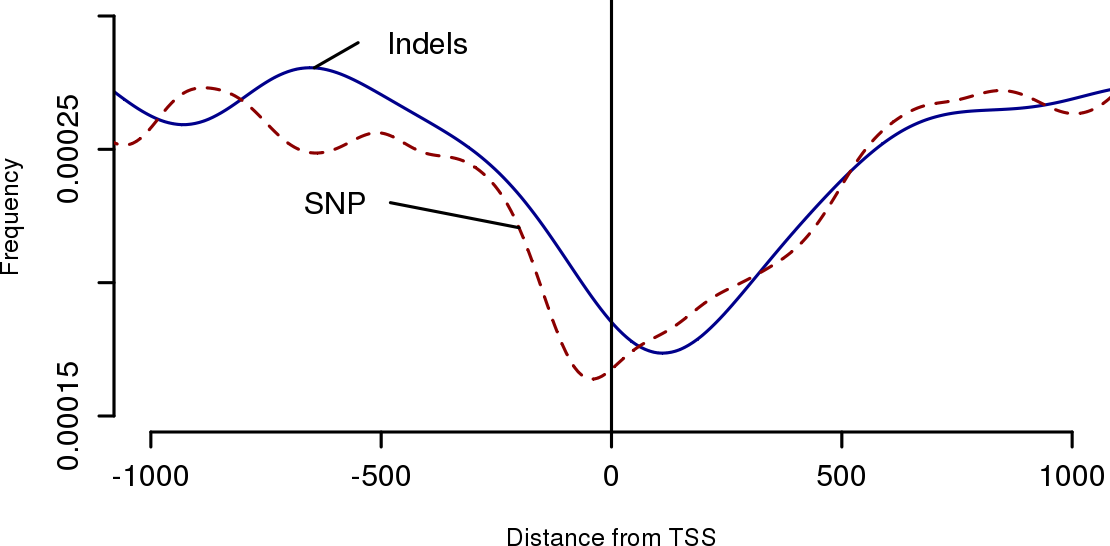
Density distribution of SNPs and Indels in promoters of the human cell line GM12878.

